# Intratumoral administration of recombinant murine interleukin-12 prevents tumor progression and bone invasion

**DOI:** 10.1101/2023.05.26.542415

**Authors:** Shun Kasahara, Toshihiro Uchihashi, Toshihiro Inubushi, Kyoko Kurioka, Akinari Sugauchi, Kazuaki Miyagawa, Mikihiko Kogo, Susumu Tanaka

## Abstract

**Background:** Oral squamous cell carcinoma (OSCC) progression is accompanied by bone invasion. Therefore, maintaining oral function is necessary to regulate tumor progression. Also, interleukin-12 (IL-12), a well-known anti-tumor cytokine, can suppress osteoclast differentiation in vitro. Accordingly, this study evaluated the therapeutic effects of locally administered IL-12 in an immunocompetent mouse model with mandibular bone invasion mimicking clinical features.

**Methods:** We investigated anti-bone resorption effects using SCCVII subcutaneous and bone invasion models both in immunocompetent and athymic mice. Furthermore, we measured bone resorption using micro-computed tomography.

**Results:** Intratumoral injection of recombinant murine IL-12 (r-mIL-12) significantly prolonged immunocompetent mouse survival and suppressed tumor growth and bone resorption. Real-time PCR analysis revealed that interferon-gamma (IFN-γ) and Fas ligand (FasL) were upregulated after r-mIL-12 administration, compared to control levels. However, when the athymic mouse bone invasion model was evaluated, r-mIL-12-mediated suppression of tumor growth and bone resorption were equivalent to those observed in the control group, highlighting the key role of T cells in the bone invasion.

**Conclusions:** r-mIL-12 may represent a potent therapeutic agent for OSCC accompanied by bone invasion.

**SUMMARY:** Intratumoral injection of recombinant murine IL-12 showed anti-tumor and anti-bone resorption effects in an immunocompetent mouse bone invasion model through a T cell-dependent mechanism.

## INTRODUCTION

Oral squamous cell carcinoma (OSCC) is very common worldwide and its incidence has been increasing in recent years (Ghantous and Abu Elnaaj, 2017). OSCC progression drastically decreases the quality of life (QOL) of patients by impeding oral functions, leading to masticatory dysfunction, dysarthria, dysphagia, respiratory disorders, and poor aesthetics (Sharma et al., 2019). Furthermore, OSCC, particularly gingival cancer, frequently invades the adjacent mandibular bone. Since jawbone invasion is strongly associated with OSCC metastasis and poor prognosis (Michalek et al., 2019; Okura et al., 2016), the control of OSCC progression, especially jawbone invasion, is extremely important. The standard approach used to treat OSCC with bone invasion involves surgery with enlarged segmental resection, resulting in a considerable decrease in QOL for patients (Deshmukh and Shekar, 2021). Similarly, radiotherapy, with or without chemotherapy, can cause tissue deficits and subsequent osteoradionecrosis, which can be difficult to treat (Minhas et al., 2017).

The pleiotropic cytokine interleukin-12 (IL-12) plays an important role in anti-tumor immunity (Tugues et al., 2015). For instance, IL-12 mediates the shift in the differentiation of CD4+ T-helper (Th)1 cells from CD4+ Th0 cells (Vacaflores et al., 2016). It also stimulates cytotoxic T cells (CTLs) and natural killer (NK) cells (Andersen et al., 2006; Markiewicz et al., 2009). Owing to its ability to activate both adaptive (CTLs) and innate (NK cells) immunities, IL-12 is one of the most promising candidates for tumor therapy (Lasek et al., 2014). Besides the potent effects of IL-12 in anti-tumor immunity, it can suppress osteoclast formation by increasing the expression of interferon-gamma (IFN-γ) (Amarasekara et al., 2018; Gillespie, 2007; Nagata et al., 2003). In addition, IL-12 induces apoptotic changes in osteoclasts by increasing Fas ligand (FasL) expression (Kitaura et al., 2002; Yoshimatsu et al., 2009). As described above, the prevention of bone invasion is important for the control of OSCC progression. Since bone invasion in carcinoma is caused by activated osteoclasts in the tumor microenvironment (TME) rather than directly by the carcinoma itself (Jimi et al., 2011; Semba et al., 1996), we hypothesized that IL-12 may inhibit bone destruction in OSCC indirectly. While clinical trials attempting treatment with systemic administration of IL-12 have often failed to owe to toxic side effects (Cohen, 1995; Leonard et al., 1997; Wang et al., 2017). However, local IL-12 application appears to present a promising safety profile with similar or better anti-tumor efficacy than that of systemic administration (Nguyen et al., 2020). Therefore, an effective local IL-12 application may represent a useful approach for OSCC accompanied by jawbone invasion due to its anti-bone resorption and anti-tumor efficacy. However, parallel IL- 12 anti-tumor and anti-bone resorption efficacy have not been investigated previously.

Thus, this study investigated the anti-tumor effects and underlying mechanisms of actions of locally administered recombinant murine IL-12 (r-mIL-12) in OSCC mouse models showing mandibular bone invasion (Nomura et al., 2007).

## RESULTS

### r-mIL-12 shows no cytotoxicity in SCC□ cells in vitro

To evaluate any possible direct cytopathic effects of r-mIL-12 other than effects exerted via an immune pathway, the anti-tumor effects of r-mIL-12 were evaluated in vitro. There was no significant difference in the number of SCC□ cells between the control and r-mIL-12 groups. This indicates that r-mIL-12 treatment has no direct cytopathic effects on SCC□ cells in vitro (Fig. 1).

**Fig. 1.**
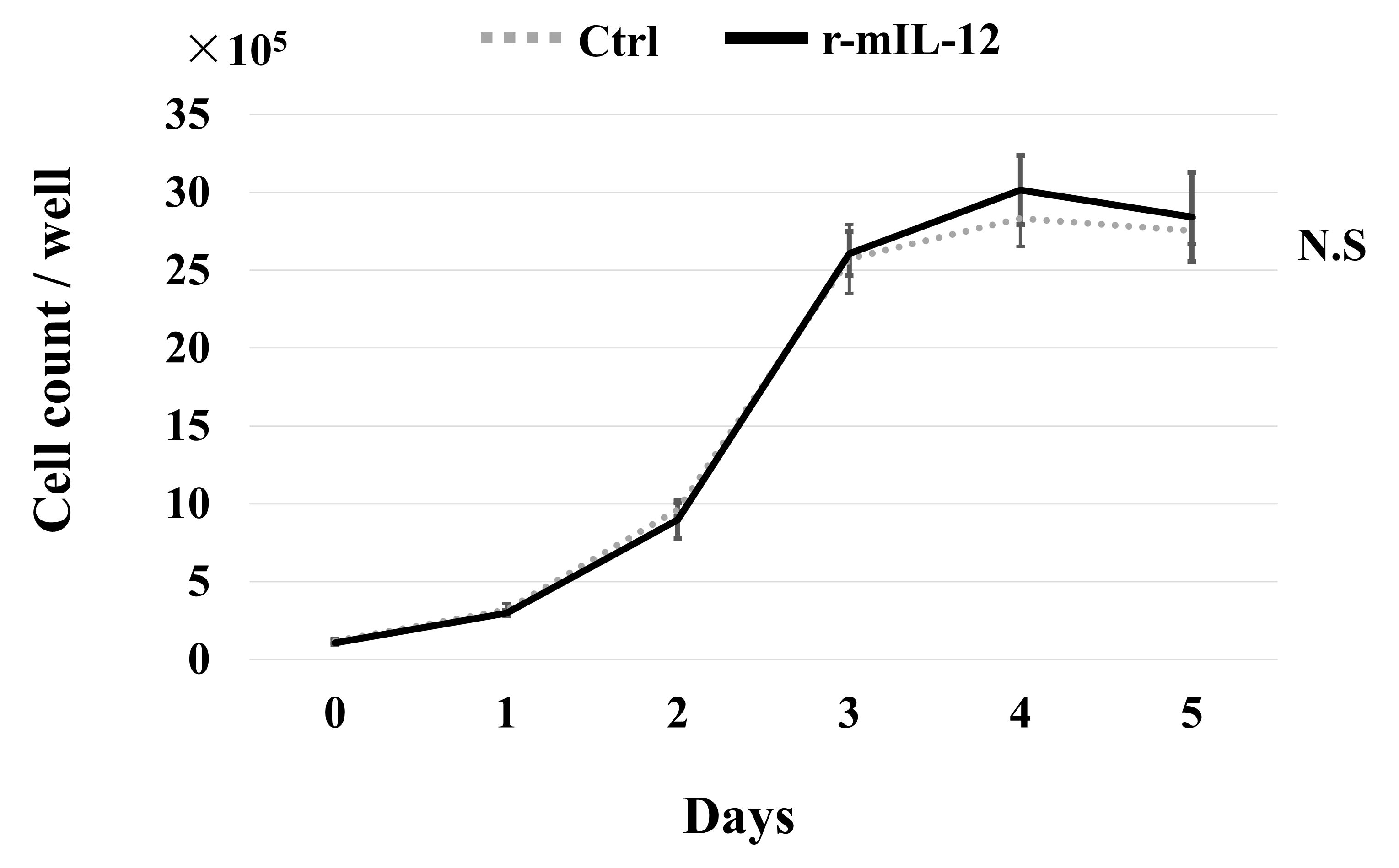
In vitro cytotoxicity testing of r-mIL-12. SCC□ cells were seeded into 6-well plates and cultured in a medium containing r-mIL-12 or PBS as control. Surviving cells were counted daily. The results show that r-mIL-12 was not cytotoxic in cultured SCC□ cells. Bars indicate the mean ± standard error of the mean. N.S., not significant (Student’s t-test). r-mIL-12, recombinant murine interleukin-12; PBS, phosphate-buffered saline.

### r-mIL-12 shows anti-tumor efficacy in an immunocompetent mouse subcutaneous tumor model

To examine the anti-tumor effects of locally administered r-mIL-12, we injected r-mIL-12 intratumorally in an immunocompetent mouse subcutaneous tumor model containing SCC□ cells. The experimental timeline is shown in Figure 2A. Intratumoral injection of r-mIL-12 significantly suppressed the growth of subcutaneous tumors (Fig. 2B) and prolonged overall survival (Fig. 2C). While the subcutaneous model is ideal for studying anti-tumor effects against SCCVII cells in vivo, the effects of r-mIL-12 on bone invasion cannot be evaluated with it. As mentioned above, bone invasion in carcinoma is caused by the activation of osteoclasts in the TME rather than being a direct effect of the carcinoma. To investigate the inhibitory effect of r-mIL-12 injection on bone invasion, we used an SCCVII bone invasion model that mimicked the behavior of mandibular gingival SCC in clinical settings (Fig. 3A). Injection of r-mIL- 12 significantly suppressed tumor growth (Fig. 3B) and tended to prolong overall survival (Fig. 3C). These results indicate that intratumoral administration of r-mIL-12 inhibits tumor growth in vivo in both the subcutaneous model and the bone invasion model.

**Fig. 2.**
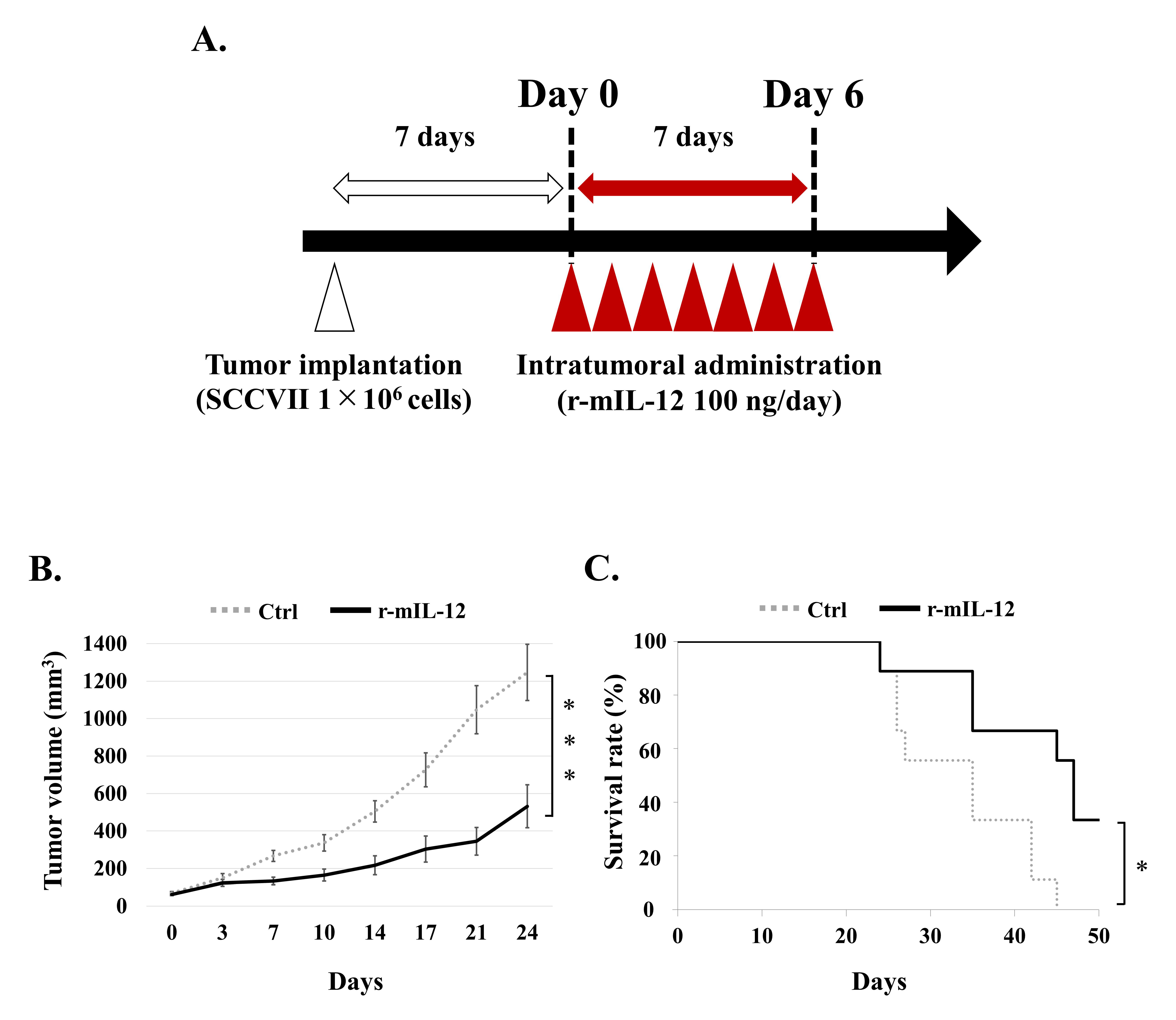
Anti-tumor effects of r-mIL-12 in an immunocompetent mouse subcutaneous tumor model. **A.** The experimental schedule was as follows: Day 0 was defined as 7 d after tumor implantations (1 × 10^6^ SCCVII cells). r-mIL-12 (2 ng/µL/d) was administered into the subcutaneous tumor for seven consecutive days (days 0–6) and the tumor volume was measured three times per week. Mice were euthanized when the tumor reached 20 mm in diameter. **B, C.** r-mIL-12 significantly suppressed the growth of subcutaneous tumors (B) and prolonged overall survival (C). Tumor volume (mm^3^) = length × width × height (mm). Control, N = 10; r-mIL-12, N = 9. Bars, mean ± standard error. ***, *P* < 0.001 (Student’s t-test); *, *P* < 0.05 (Log-rank test). r-mIL-12, recombinant murine interleukin-12.

**Fig. 3.**
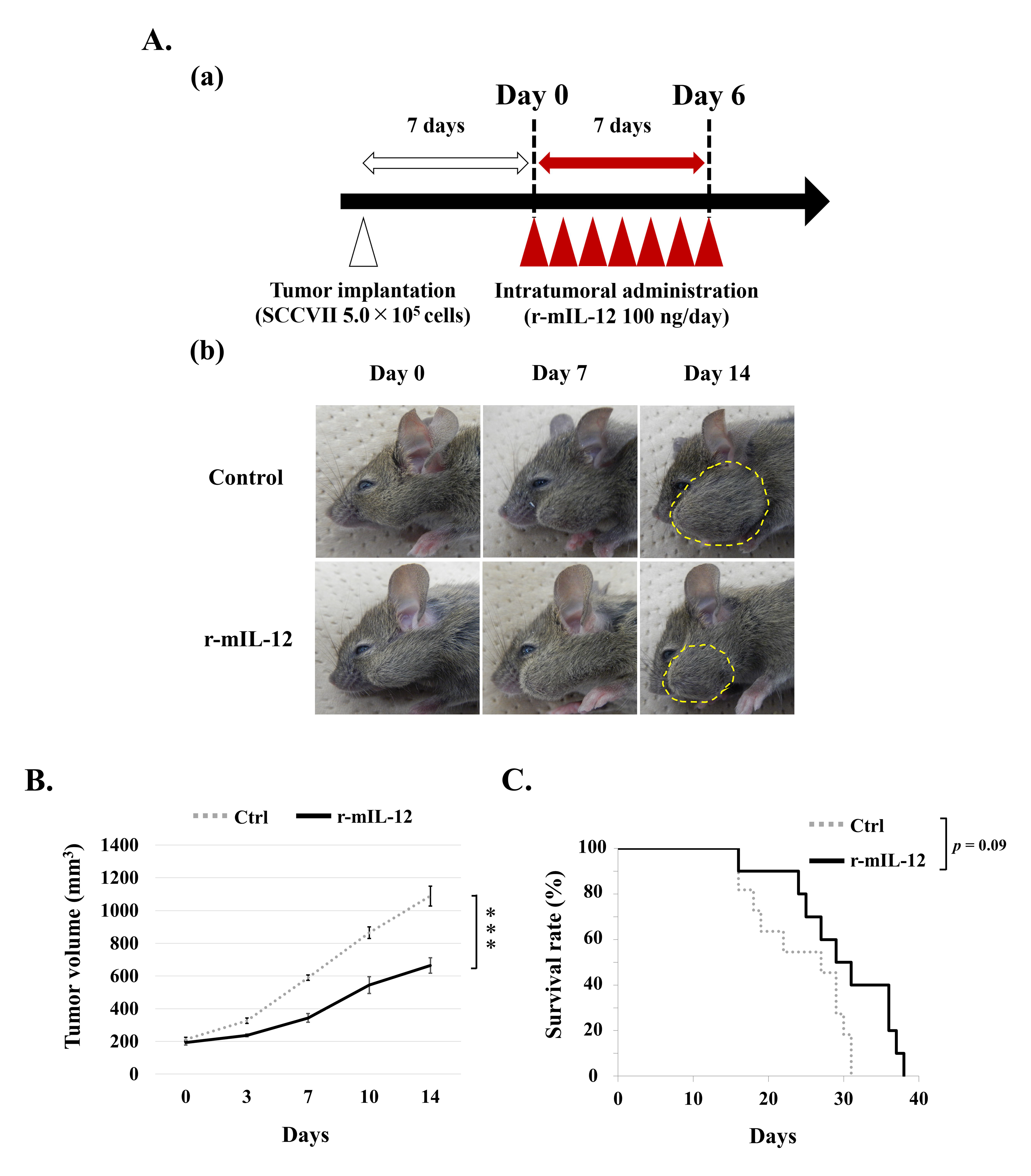
Anti-tumor effects of r-mIL-12 in an immunocompetent mouse bone invasion model. **A.** Experimental schedule (a) and representative appearance of mice at designated days (b). Day 0 was defined as 7 d after tumor implantations (5 × 10^5^ SCCVII cells). r-mIL-12 (2 ng/µL/day) was administered into the mandibular tumor for seven consecutive days (days 0–6) and the tumor volume was measured three times per week. The dashed yellow circumferences indicate the mandibular tumors. **B, C.** r-mIL- 12 administration significantly suppressed tumor growth (B) and tended to prolong overall survival (C). Tumor volume (mm^3^) = length × width × height (mm). Control, N = 11; r-mIL-12, N=10. Bars, mean ± standard error. ***, *P* < 0.001 (Student’s t-test); *P* = 0.09 (Log-rank test). r-mIL-12, recombinant murine interleukin-12.

### r-mIL-12 injection shows anti-bone resorption effects in an immunocompetent mouse bone invasion model

IL-12 is known to show anti-osteoclastogenic effects in vitro (Gillespie, 2007; Kitaura et al., 2002; Yoshimatsu et al., 2009). Thus, the anti-bone resorption effects of r-mIL-12 were investigated in an immunocompetent mouse bone invasion model. The volume of bone resorption was measured using micro-computed tomography (micro-CT) (Fig. 4A). Injection of r-mIL-12 led to significant suppression of bone resorption in the tumor group compared to that in the control group (Fig. 4B). In addition, deposition of new bone tissue on the periosteal surface near the tumor was observed before bone resorption, as it occurs frequently in clinical cases, indicating that this model reflects the clinical features of OSCC accompanied by bone invasion (Fig. 4B).

**Fig. 4.**
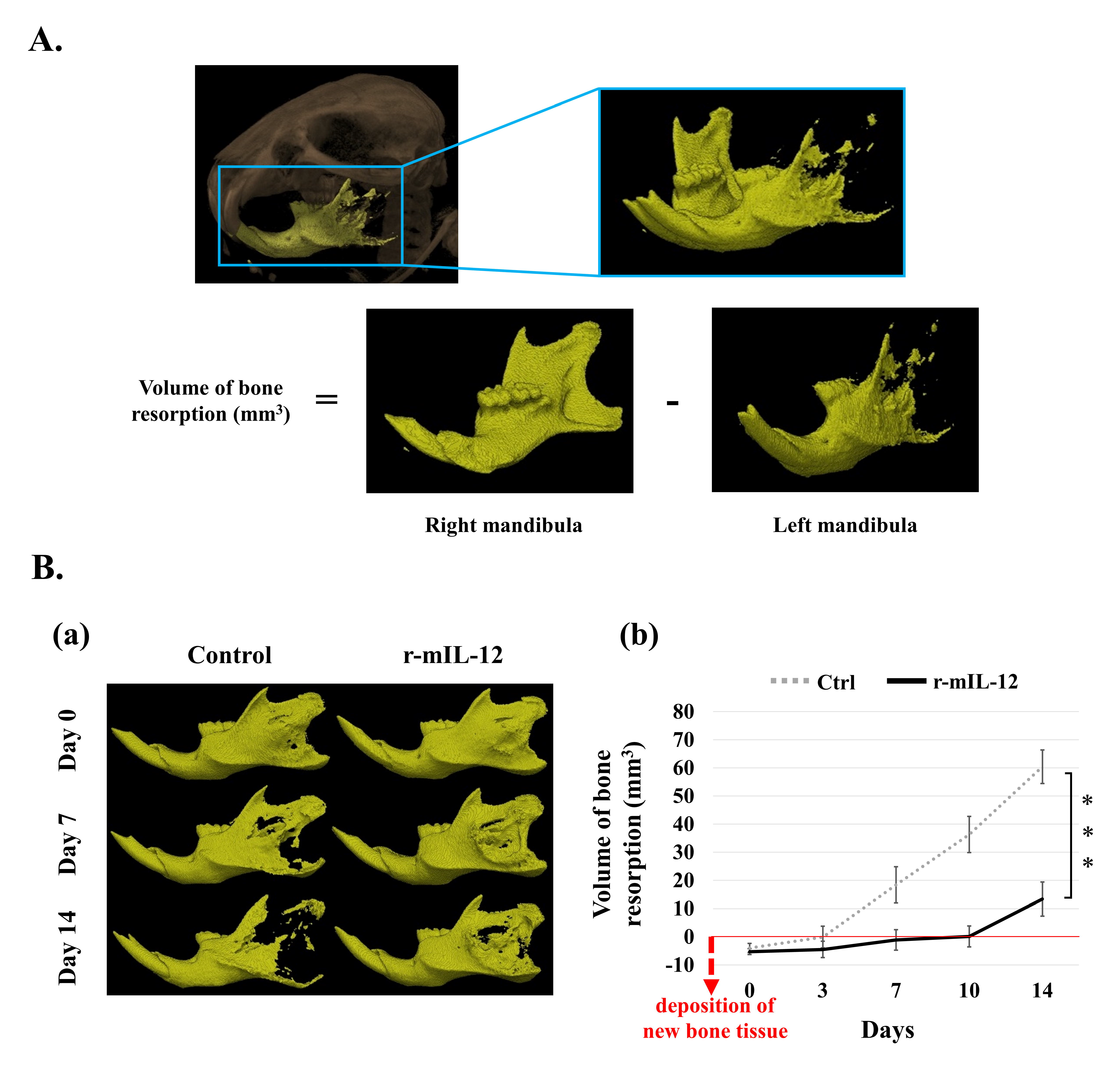
Anti-bone resorption effect of r-mIL-12 in a mouse bone invasion model. **A.** Method of measuring the volume of bone resorption via micro-CT. The mandibular bone was extracted as indicated by the blue area. The volume of bone resorption was defined as follows: volume of bone resorption = volume of the right mandibular - volume of the left mandibula. **B. (a)** The volume of bone resorption was measured via micro-CT. Representative images from days 1, 7, and 14 are shown. **(b)** r-mIL-12 injection showed significant suppression of bone resorption compared to that obtained using control injection. Deposition of new bone tissue on the periosteal surface was observed prior to bone resorption. The red line indicates the baseline. The dashed red arrowhead indicates the deposition of new bone tissue. Control, N = 9; r-mIL-12, N = 8. Bars, mean ± standard error. ***, *P* < 0.001 (Student’s t-test). r-mIL-12, recombinant murine interleukin-12; micro-CT, micro-computed tomography.

### r-mIL-12 significantly decreases the number of osteoclasts in histomorphometric analyses

We performed hematoxylin and eosin (H-E) staining as well as immunohistochemical staining for cathepsin K, an osteoclast marker, and investigated the anti-bone resorption effect of r-mIL-12 by calculating the number of osteoclasts/bone perimeter (N.Oc/B.Pm) and osteoclast surface/bone surface (Oc.S/BS) as histomorphometric analysis (Fig. 5A). The results showed that Oc.S/BS and N.Oc/B.Pm significantly decreased following intratumoral administration of r-mIL-12 compared to those of the control (Fig. 5B). These results suggest that r-mIL-12 may demonstrate its anti-bone resorption effect by inhibiting the induction of osteoclasts.

**Fig. 5.**
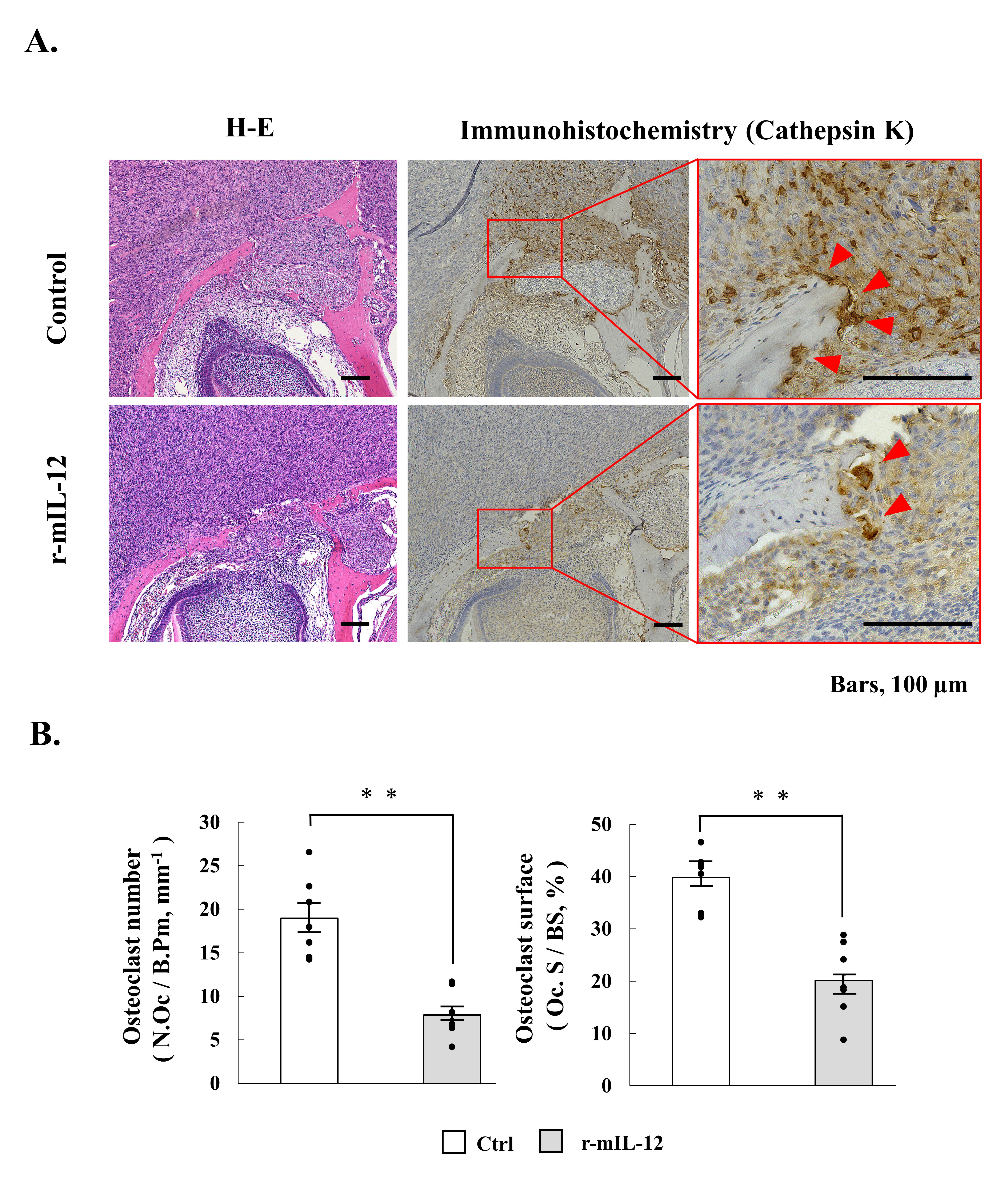
Histological analysis. **A.** The bone invasion model mice were euthanized on day 7, after which H-E staining and cathepsin K immunohistochemistry was performed. **B.** Histomorphometric analyses showed that osteoclast number (the number of osteoclasts/bone perimeter [N.Oc/B.Pm]) and osteoclast surface (osteoclast surface/bone surface [Oc.S/BS]) significantly decreased following intratumoral administration of r-mIL-12. Scale Bar, 100 μm. N = 7 in each group. Bars, mean ± standard error. **, *P* < 0.01 (Mann–Whitney U test). H-E, hematoxylin, and eosin; r-mIL-12, recombinant murine interleukin-12.

### r-mIL-12 increases the number of CD3, CD4, and CD8 cells in mandibular tumors

To further investigate the influence of r-mIL-12 on immune cells in the TME, we performed immunohistochemical staining in r-mIL-12-injected regions. Immunohistochemical staining showed that r-mIL-12 administration increased the number of CD3+, CD4+, and CD8+ T cells in the vicinity of the mandibular bone where r-mIL-12 had been injected, compared to that where phosphate-buffered saline (PBS) had been injected (Fig. 6A). These results indicate that r-mIL-12 injection caused accumulation of T cells in r-mIL-12-injected cancer lesions. It also suggests that the increase in anti-tumor immunity following r-mIL-12 injection significantly inhibited tumor growth in the model.

**Fig. 6.**
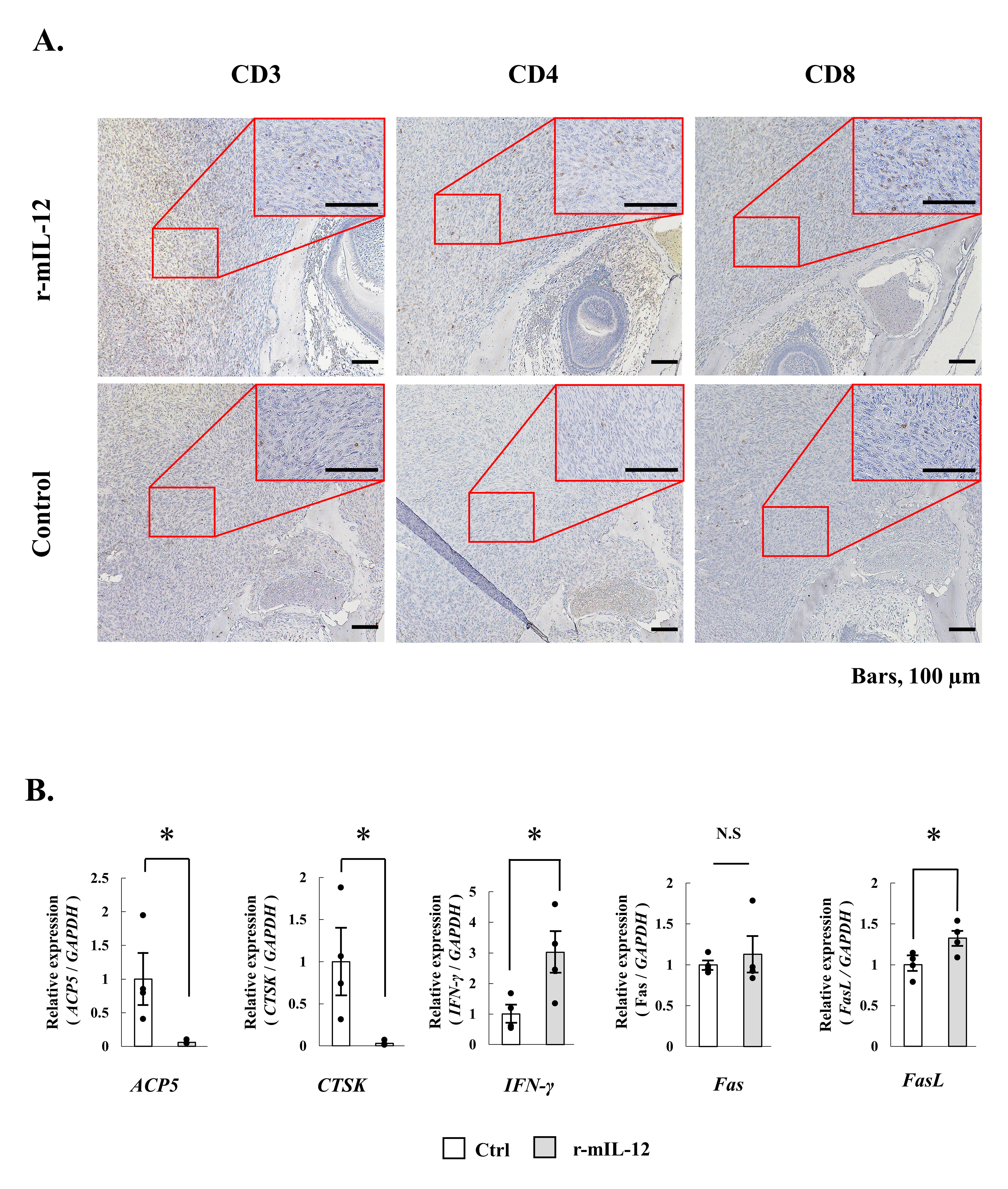
Effect of r-mIL-12 administration on the number of T cells and gene expression in mandibular tumors. **A.** Bone invasion model mice were euthanized on day 7 and immunohistochemistry analyses were performed, showing that r-mIL-12 administration increased the numbers of CD3+, CD4+, and CD8+ T cells in mandibular tumors. Scale Bar, 100 μm. **B.** A portion of the mandibular tumor was extracted from the bone invasion models on day 7 and mRNA expression levels of *Acp5*, *Ctsk*, *Ifng*, *Fas*, and *Fasl* were determined via qPCR. *Ifng* and *Fasl*, which are closely associated with T cells, were significantly upregulated following r-mIL-12 administration. However, r-mIL-12 administration downregulated *Acp5* and *Ctsk*, which are known to be osteoclast markers. N = 4 in each group. Bars, mean ± standard error. *, *P* < 0.01 (Mann–Whitney U test). N.S., not significant. r-mIL-12, recombinant murine interleukin-12; qPCR, quantitative PCR; *Acp5,* acid phosphatase 5; *Ctsk,* cathepsin K; *Ifng,* interferon gamma.

### r-mIL-12 significantly decreases the expression of Acp5 and Ctsk accompanied by significant increases in IFN-γ and Fasl expression

Quantitative real-time PCR analysis results showed that r-mIL-12 administration significantly upregulated *Ifng* and *FasL* (Fig. 6B), which are associated with the activation of T cells. This was consistent with the accumulation of T cells observed in immunohistochemistry analyses. Further, r-mIL-12 administration significantly downregulated *Acp5* and *Ctsk* (Fig. 6B), which are known to be osteoclast markers. There was no statistical difference in *Fas* expression between the treatment and control groups. The findings revealed that T cell activation induced by r-mIL-12 injection promoted *Ifng* and *Fasl* expression accompanied by downregulation of *Acp5* and *Ctsk*, suggesting that r-mIL-12 (i) inhibited differentiation of pre-osteoclasts to osteoclasts via increased Fas-FasL pathway activation and/or (ii) suppressed osteoclast formation via increased production of IFN-γ, thereby exerting anti-tumor immune effects via an IFN-γ pathway.

### r-mIL-12 does not show additional anti-tumor and anti-bone resorption effects in an athymic mouse bone invasion model

Based on the observation that r-mIL-12 significantly downregulated *Acp5* and *Ctsk* as well as significantly upregulated *Ifng* and *Fasl* in an immunocompetent mouse model, we assumed that r-mIL-12 exerted its anti-tumor and anti-bone resorption effects via T cells. To clarify the involvement of T cells in the inhibitory effects of r-mIL-12 on bone invasion, we used an athymic mouse bone invasion model (Nomura et al., 2007), measured the tumor volume, and analyzed bone resorption levels through micro-CT analysis. Although deposition of new bone tissue was observed before bone resorption, as indicated in immunocompetent mice, our data showed that both anti-tumor (Fig. 7A) and anti-bone resorption effects (Fig. 7B) arising from r-mIL-12 were attenuated in athymic mice to almost the same levels as in the control. In addition, real-time PCR analysis revealed that *Acp5* and *Ctsk* expression following injection of r-mIL-12 was equivalent to that in the control, indicating that osteoclast numbers were not influenced by r-mIL-12 injection. However, *Ifng* and *Fasl* expression still increased significantly following the administration of r-mIL-12 (Fig. 7C). In this model, mandibular tumors grew rapidly in mice injected with either a control solution or r-mIL-12, suggesting that IFN-γ was not involved in anti-tumor effects, with secretion possibly occurring from other lymphocytes such as NK cells. Furthermore, cells in the TME can express FasL, which is known to lead to apoptosis in tumor-infiltrating lymphocytes (Zhu et al., 2019). The micro-CT results in this model indicate that the high expression of FasL was irrelevant to the apoptotic change of osteoclasts. Furthermore, they strongly suggest that the anti-bone resorption effect of r-mIL-12 in SCCVII mouse models utilizes a T cell-dependent mechanism.

**Fig. 7.**
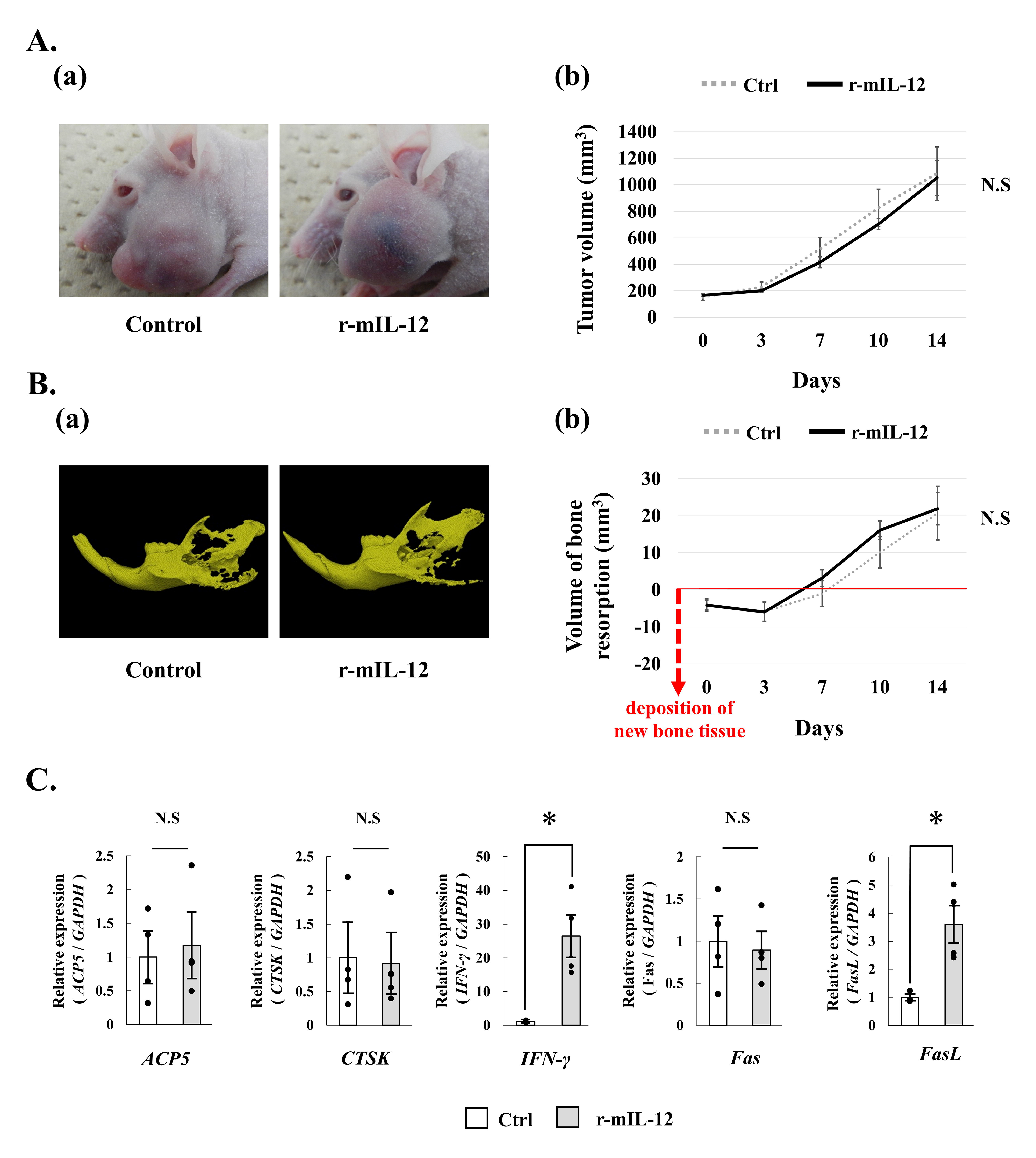
Therapeutic effect of r-mIL-12 in an athymic bone invasion mouse model. **A, B.** In athymic mice, the anti-tumor (A) and anti-bone resorption effects (B) of r-mIL- 12 were reduced to almost the same levels as those in controls. **A (a).** Representative photos of mice at day 14 injected with control or r-mIL-12. **(b).** The tumor volumes injected with r-mIL-12 or PBS were similar. **B (a).** Representative image of the mandibular bone at day 14 injected with control or r-mIL-12. **(b).** Anti-bone resorption effects of r-mIL-12 were attenuated compared to those of the control. **C.** Real-time PCR analysis showed that *Acp5, Ctsk, and Fas* expression levels were similar following the injection of r-mIL-12 or control. However, *Ifng* and *Fasl* expression increased significantly following injection of r-mIL-12, compared to the control. Ctrl, N = 5; r-mIL-12, N = 4. Tumor volume (mm^3^) = length × width × height (mm). Bars, mean ± standard error. N.S., not significant (Mann-Whitney U test). r-mIL-12, recombinant murine interleukin-12.

## DISCUSSION

OSCC with jawbone invasion is classified as T4a, staged into IV (Deshmukh and Shekar, 2021; Okura et al., 2016), and standard therapy involves surgical resection with jawbone excision, leading to severe degradation in patients’ QOL. Jawbone invasion is a prognostic factor mediated by osteoclast activation (Deshmukh and Shekar, 2021; Michalek et al., 2019), not by tumor-mediated direct effects, as reported previously (Jimi et al., 2011; Semba et al., 1996). Building on the known anti-bone resorption effects of IL-12 against OSCC, we aimed to determine whether local IL-12 administration could represent a minimally invasive treatment that would regulate bone resorption occurring resulting from tumor development. We investigated anti-bone resorption effects using an SCCVII subcutaneous bone invasion model both in immunocompetent and athymic mice. In the immunocompetent SCCVII subcutaneous bone invasion model, tumor growth was significantly inhibited following r-mIL-12 injection, whereas in the athymic mouse model, r-mIL-12 did not show tumor inhibition effects greater than those of the control. This indicates that T cells play an important role in the anti-cancer effects exerted by IL-12. Histological analysis showed increased accumulation of CD3+, CD4+, and CD8+ T cells following r-mIL-12 injection, which further supports the importance of T cells in local IL-12 therapy.

In this study, we measured bone resorption using micro-CT, which enabled us to measure the volume of bone destruction as a function of time. Remarkably, we observed the deposition of new bone tissue on the periosteal surface near the tumor prior to bone destruction. This morphological change is frequently observed in clinical cases (Semba et al., 1996; Totsuka et al., 1991), indicating that our model accurately mimicked the clinical course of OSCC with bone invasion. Micro-CT analysis results revealed significant anti-bone resorption effects in the immunocompetent mouse model. However, there was no significant difference in bone changes between the r-mIL-12 and control groups in the athymic mouse model. Immunohistological analysis results showed that treatment with r-mIL-12 significantly decreased the number of cathepsin K-positive multinuclear cells on periosteal surfaces in the immunocompetent mouse model. The results of real-time PCR showed significant downregulation of *Acp5* and *Ctsk* only in the immunocompetent mouse model, and not the athymic mouse model, which was consistent with the decrease in osteoclast numbers in the immunocompetent mouse model. Further, r-mIL-12 administration significantly upregulated *Ifng* as well as *Fasl*, known to be largely expressed by T cells or tumor cells (Andersen et al., 2006). Considering that r-mIL-12 significantly inhibited tumor growth only in the immunocompetent mouse model, a high *Fasl* expression in this experiment appeared to be influenced by T cells. These results suggest that r-mIL-12 suppresses osteoclast formation by increasing IFN-γ expression and induces apoptotic changes in osteoclasts by increasing FasL expression in T cells, thereby suppressing mandibular bone resorption.

Accumulation of CD3+, CD4+, and CD8+ T cells after r-mIL-12 injection was observed adjacent to the mandibular periosteum region where r-mIL-12 was injected, while a lower level of accumulation of those cells was observed in the tumor central region (data not shown), suggesting that r-mIL-12 administered locally remains in the vicinity of the injection site and exerts its anti-tumor effect locally. This phenomenon suggests that local injection of r-mIL-12 may have better safety characteristics and may avoid the possibility of a lethal cytokine storm.

Several strategies for IL-12 administration have been explored to overcome the problems of toxicity, and several localized IL-12 delivery approaches are now being tested clinically (Nguyen et al., 2020). Most recently, oncolytic viruses that directly lyse tumor cells and express active IL-12 only within the TME have been explored as potential treatments (Alkayyal et al., 2016; Nguyen et al., 2020). Triple-mutated oncolytic herpes virus G47Δ is a promising agent for various solid cancers and was approved in Japan as a new drug for malignant glioma in 2021 (Fukuhara et al., 2021). A recent report indicated that G47Δ represents a potent candidate agent for OSCC, particularly in cases with lymph node metastasis (Uchihashi et al., 2021). In addition, a bacterial artificial chromosome system that allows the addition of a gene of choice within the G47Δ backbone may allow efficient regulation of bone invasion if an IL-12 gene is inserted, leading to continuous expression during infection (Fukuhara et al., 2005).

IL-12 exerts a potent effect via T cells, and this effect may be enhanced in combination treatments with immune checkpoint inhibitors (ICIs), which activate T cells. Treatment with ICIs such as anti-PD1 antibody pembrolizumab and anti-CTLA-4 antibody nivolumab in combination with IL-12 has shown potential therapeutic effects in clinical trials (Algazi et al., 2020; Berraondo et al., 2018; Chiocca et al., 2022).

In summary, local IL-12 administration affecting T cells appears to represent a suitable and efficacious candidate therapy for OSCC with bone invasion in clinical settings. Potential combination therapy with recent immunotherapy and/or gene therapy agents might represent an even more promising treatment for OSCC.

## MATERIALS AND METHODS

### Cell culture

The C3H/He mouse squamous cell carcinoma cell line SCC□ was provided by Professor Eijiro Jimi, Kyushu University. SCC□ cells were maintained in Dulbecco’s modified Eagle’s medium (Thermo Fisher Scientific, Waltham, MA, USA) supplemented with 10% (v/v) fetal bovine serum (Thermo Fisher Scientific) and incubated at 37 °C with 5% CO_2_.

### Animal studies

Female C3H/He and BALB/c *nu/nu* mice (age, 5–6 weeks) were obtained from SLC (Shizuoka, Japan). All experimental animal protocols were approved by the Committee for Ethics of Animal Experimentation and were performed following the Guidelines for Animal Experiments at Osaka University (28-003-0). Adequate measures were taken to always minimize animal pain and discomfort.

### Cytopathic effects of IL-12

r-mIL-12 was obtained from R&D Systems (Minneapolis, MN, USA) and diluted with PBS to 2 ng/µL. SCC□ cells were seeded into 6-well plates (Corning, Corning, NY, USA) and cultured in a medium containing r-mIL-12 (5 µL/well) or PBS (5 µL/well) as control. The number of surviving cells was counted daily using a Countess II FL Automated Cell Counter (Thermo Fisher Scientific).

### Anti-tumor effect of r-mIL-12 on in vivo mouse models

To prepare subcutaneous tumor models, SCCVII cells (1.0 × 10^6^ cells/50 µL) were injected subcutaneously into the left flank of C3H/He mice under general anesthesia. To prepare bone invasion models, SCCVII cells (5.0 × 10^5^ cells/50 µL) were injected into the central region of the left masseter of C3H/He or BALB/c-*nu/nu* mice. Day 0 was defined as 7 d after SCC□ cell injection. To test the anti-tumor effects, r-mIL-12 (2 ng/µL/d) was administered into the formed mass adjacent to the mandibular periosteum over seven consecutive days (days 0–6). As a control, PBS (50 µL) was injected into the tumor on the same schedule. Tumor volume was measured three times per week. Mice were euthanized when the tumor reached 20 mm in diameter.

### Anti-bone resorption effect of r-mIL-12 in mouse bone invasion model

To test the anti-bone resorptive effects, r-mIL-12 (2 ng/µL/d) was administered to tumors in mouse bone invasion models for seven consecutive days (days 0–6). The control group was injected with 50 µL of PBS on the same schedule, and the tumor and bone resorption volumes were compared.

Bone resorption was measured using micro-CT (R-mCT2; Rigaku, Tokyo, Japan) on days 0, 3, 7, 10, and 14. The volume of bone resorption was defined as (volume of the right mandibula) - (volume of the left mandibula). The operating conditions for R-mCT2 were as follows: tube voltage, 90 kV; tube current, 200 µA; field of view, 20 mm. The data were analyzed using TRI /3 D-Bon64 (RATOC, Tokyo, Japan).

### Histological analysis

The bone invasion model mice were euthanized on day 7. Collected tissues were fixed with 10% (v/v) formalin neutral buffer solution (FUJIFILM Wako Pure Chemical Corporation, Osaka, Japan) for 72 h at room temperature (25–28 °C) and then, decalcified using 15% (v/v) EDTA for 5 weeks. After decalcification, samples were embedded in paraffin and sliced into 4–5 µm sections using a microtome. Immunohistochemical staining was performed as follows: the primary antibodies used included anti-cathepsin K (sc-48353, 1:200 dilution; Santa Cruz Biotechnology, Dallas, TX, USA), anti-mouse CD3 (ab16669, 1:250 dilution; Abcam, Cambridge, United Kingdom), anti-CD4 (ab183685, 1:1000 dilution; Abcam), and anti-CD8 (ab217344, 1:2000 dilution; Abcam). A VECTASTAIN ABC Kit (Vector Laboratories, Burlingame, CA, USA) was used to detect secondary antibodies. Immunostaining was completed using a DAB Substrate Kit (Vector Laboratories). Also, a small number of sections were stained with H-E.

The osteoclast number (N.Oc/B.Pm) and osteoclast surface (Oc.S/BS) were evaluated in histomorphometric analyses (Oikonomidou et al., 2016). N.Oc/B.Pm was defined as the number of cathepsin K-positive multinuclear cells on the bone surface in contact with the tumor. Oc.S/BS was defined as the length of the bone surface covered with cathepsin K-positive multinuclear cells.

### Real-time PCR analysis

A portion of the tumor adjacent to the mandibular periosteum of bone invasion models was extracted on the day 7, and total RNA was isolated using the RNeasy Mini Kit (QIAGEN, Germantown, MD, USA). cDNA was synthesized using 2 µg total RNA using a High-Capacity cDNA Reverse Transcription Kit (Applied Biosystems, Foster City, CA, USA). Real-time PCR was performed using a StepOnePlus system (Applied Biosystems) with a TaqMan Gene Expression Assay (20×) (Applied Biosystems) to amplify the following genes: acid phosphatase 5 (*Acp5*) (Mm00438864m1), cathepsin K (*Ctsk*) (Mm00484039m1), *Ifng* (Mm01168134m1), *Fas* (Mm01204974m1), *Fasl* (Mm00438864m1), and glyceraldehyde 3-phosphate dehydrogenase (*Gapdh*) (Mm99999915g1). Real-time PCR was performed using a 10 µL mixture of 4.5 µL cDNA template, 5 µL TaqMan Universal PCR Master Mix (2×) (Applied Biosystems), and 0.5 µL of each TaqMan Gene Expression Assay reagent. mRNA Ct values for these genes were normalized using those of *Gapdh* (Mm99999915g1) and expressed as increases or decreases relative to the values of the control group.

### Statistical analysis

The Student’s *t*-test was used to analyze data from the r-mIL-12 cytotoxicity assay, tumor volume, and bone resorption volume. Real-time PCR data were analyzed using the Mann–Whitney U test. The Kaplan–Meier analysis was used for survival studies, and the significance was evaluated using the log-rank test. Values of *P* < 0.05 indicated significance.

## Acknowledgments

We thank Professor Eijiro Jimi for providing the C3H/He mouse squamous cell carcinoma cell line SCC□.

## Author contributions

Conceptualization: T.U., T.I., M.K., S. T; Methodology: S.K., T.U.; Investigation: S.K., T.U., T.I., A.S., K.K., K.M.; Resources: T.U., S.T.; Data curation: A.S., K.K., K.M.; Writing – original draft: S.K., T.U.; Writing – review & editing: T.U., T.I.; Supervision: M.K., S.T.; Project administration: T.U.; Funding acquisition: T.U.

